# NLSDeconv: an efficient cell-type deconvolution method for spatial transcriptomics data

**DOI:** 10.1101/2024.06.13.598922

**Authors:** Yunlu Chen, Feng Ruan, Ji-Ping Wang

## Abstract

**Summary:** Spatial transcriptomics (ST) allows gene expression profiling within intact tissue samples but lacks single-cell resolution. This necessitates computational deconvolution methods to estimate the contributions of distinct cell types. This paper introduces NLSDeconv, a novel cell-type deconvolution method based on non-negative least squares, along with an accompanying Python package. Benchmarking against 18 existing deconvolution methods on various ST datasets demonstrates NLSDeconv’s competitive statistical performance and superior computational efficiency.

**Availability and implementation:** NLSDeconv is freely available with tutorial at https://github.com/tinachentc/NLSDeconv as a Python package.

## 1 Introduction

Spatial transcriptomics (ST), crowned the Method of the Year in 2020 (Marx, 2021), has revolutionized biomedical research by allowing quantification of the mRNA expression of a large number of genes simultaneously in the spatial context of the tissue. The sequencing-based ST technology, while achieving its popularity for spatial profiling of gene expression across larger tissue areas with higher throughput than the image-based ST methods, lacks single-cell resolution. Consequently, the gene expression measured within a spot or a section of the tissue only reflects the average expression of a mixture of cell populations. Understanding the precise cell type composition within each spatial location is crucial for studying developmental biology and cancer biology. Thus, deconvoluting the cell type composition within each spot has become an imperative task in ST research.

Cell-type deconvolution in ST data often utilizes a set of reference gene expression profiles for known cell types, frequently obtained from scRNA-seq studies (Moses and Pachter, 2022). Many methods have been developed in recent years, including probabilistic-based methods (Andersson et al., 2020; Ma and Zhou, 2022; Kleshchevnikov et al., 2022; Lopez et al., 2022; Cable et al., 2022; Danaher et al., 2022); non-negative matrix factorization-based methods (Rodriques et al., 2019; Dong and Yuan, 2021; Elosua-Bayes et al., 2021); and deep learning-based methods (Biancalani et al., 2021), etc. For excellent reviews of this topic, we refer to Chen et al. (2022); Li et al. (2023).

In this paper, we propose a novel method named NLSDeconv and a Python package based on the non-negative least squares method for cell-type deconvolution for ST data. We demonstrate the competitive performance of NLSDeconv relative to 18 existing methods by utilizing comprehensive benchmark datasets from both image-based and sequencing-based ST technologies, as well as the compelling efficiency of NLSDeconv.

## 2 Methods

Like many ST cell-type deconvolution methods, NLSDeconv takes two input datasets.

- An ST dataset containing RNA-seq read count for each gene at each spot within the tissue, denoted by **Y** of size *n*_*s*_ *× n*_*g*_ where *n*_*s*_ and *n*_*g*_ stand for the number of spots (spatial locations) and number of genes respectively.
- A reference single-cell RNA-seq (scRNA-seq) dataset containing scRNA-seq read count for each gene within each cell, denoted by **X** of size *n*_*c*_ *× n*_*g*_ where *n*_*c*_ stand for the number of cell *prototypes*. Each row of **X** corresponds to a cell prototype, and each prototype belongs to one of *K* unique and known cell types, though multiple cell prototypes can belong to the same single cell type. Here, we assume **X** and **Y** share the same gene dimension *n*_*g*_, but note that NLSDeconv can adjust for different dimensions through gene selection preprocessing.

The output of NLSDeconv is a cell-type deconvolution matrix 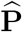 with each entry estimating the proportion of a cell *type* (not prototype) within a spot in the tissue.

The fundamental idea behind NLSDeconv is modeling the ST data’s gene read counts at a spot as a *weighted sum* of contributions from each cell prototype present. This is captured by a weight matrix **M** of size *n*_*s*_ *× n*_*c*_, where each entry indicates the weight of cell prototypes at a spot. The relationship is formulated by a linear model:

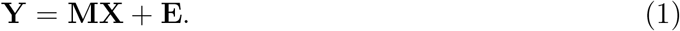

where **E** is a noise matrix of dimensions *n*_*s*_ *× n*_*g*_. This linear model captures how various cell prototypes contribute to the observed gene read counts across different spots, with **M**_*ij*_ = 0 indicating no contribution from the *j*^*th*^ cell prototype at the *i*^*th*^ spot.

In the field of ST cell-deconvolution, many existing linear models, such as those documented by Rodriques et al. (2019); Dong and Yuan (2021); Elosua-Bayes et al. (2021), typically regress **Y** onto the cell type. In contrast, our approach regresses **Y** onto the cell prototype **X**. Our method (see below) essentially builds on top of least squares objectives to solve the linear model, which differs significantly from the method in (Biancalani et al., 2021). That method employs Kullback–Leibler divergence and cosine distance to learn the matrix **M**, even though it similarly regresses **Y** on the cell prototypes **X**.

To estimate the weight matrix **M** in our method, we employ Non-negative Least Squares (NLS) (Hastie et al., 2009). Our NLS estimate 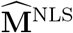 of **M** is defined by:

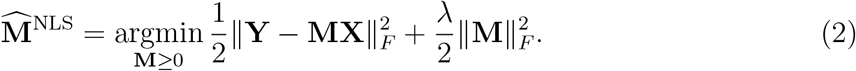

The Frobenius norm ∥*·*∥_*F*_ measures model fidelity, *λ >* 0 is a ridge regularization parameter, and the constraint **M** ≥ 0 requires all entries of the matrix **M** to be nonnegative. The NLS objective is convex, allowing projected gradient descent to converge to its global minimum (Boyd and Vandenberghe, 2004). In practice, we typically set *λ* = 0 when *n*_*s*_ ≤ *n*_*c*_ and a small positive value, *λ* = 0.1, when *n*_*s*_ *> n*_*c*_.

The method then calculates the cell-type deconvolution matrix 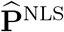 using the weight matrix estimator 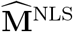. The proportion of cell type *k* ∈ {1, …, *K*} at spot *i* is calculated as follows:

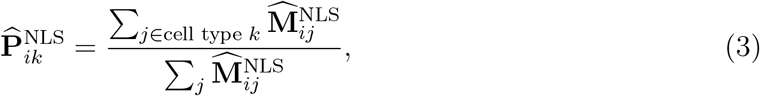

where the numerator is the sum of weights of cell type *k* at spot *i*, and the denominator is the total sum of weights across all cell types at that spot. This results in 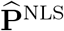, an estimator for the cell-type deconvolution matrix.

In practice, computing 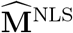, and consequentially 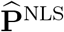 for large datasets can be challenging due to the complexities of solving the Non-negative Least Squares (NLS) minimization as described in equation (2). To enhance computational efficiency, we propose a variant of the NLS method. This approach starts with the Ordinary Least Squares (OLS) estimator:

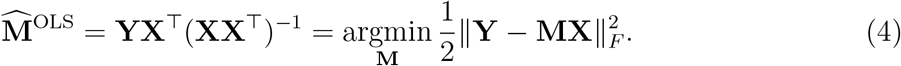

However, 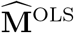 does not guarantee nonnegative entries, so we apply the soft thresholding operator, defined by (*z*)_+_ = max{*z*, 0}for *z* ∈ ℝ, to ensure nonnegativity (Hastie et al., 2009). Specifically, we estimate the proportion of cell type *k* at a spot *i* by (cf. equation (3))

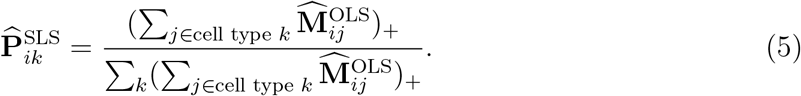

The soft thresholding ensures all entries of the cell-type deconvolution matrix 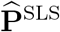 to be nonnegative.

In conclusion, we have developed two distinct cell-type deconvolution matrix estimators, 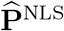 and 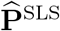. These two methods offer complementary approaches that are tailored to different computational budgets. In Section 4 below, we report our findings on the computational and statistical performance of these two estimators on benchmark ST datasets.

## 3 Software

We have developed a Python package NLSDeconv, which takes input data of all acceptable formats through *scanpy*.*read* function, e.g. h5ad, csv, text, h5, etc. The input data contains (i) the scRNA-seq reference data file, including the read count and metadata with a key indicating cell type, and (ii) the ST data file, including the read count and metadata with spatial location x and y coordinates as columns.

### Preprocess

NLSDeconv provides a data preprocessing function step for scRNA-seq data. The class *Preprocessing()* requires scRNA-seq read count, ST read count, and the cell-type key for scRNA-seq data. It performs the three ordered steps: (i) normalize each cell by total counts over all genes (*cellcount norm*) (default option is True), and (ii) remove cells with cell types observed less than a number (*cellcount min*) (default number is 2), (iii) select a certain number of genes that can most characterize each cell type using differential expression analysis (*gene top*) (default number is 200). The outputs are the preprocessed ST data and scRNA-seq data.

### Deconvolution

Two functions of cell-type deconvolution are provided. The class *Deconv()* requires scRNA-seq read count, ST read count, and the cell-type key for scRNA-seq data.

- .*SLS()* is the command for performing SLS cell-type deconvolution. The outputs are the estimated cell-type composition matrix 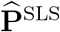, algorithm running time, and list of cell types corresponding to the column name of the resulting matrix.
- .*NLS()* is the command for performing NLS cell-type deconvolution. The required argument is learning rate (*lr*) (default is 0.1). Optional arguments are: ridge regularization parameter (*reg*), whether to use least square estimator as a warm start (*warm start*), number of epochs (*num epochs*), device for running the algorithm (*device*). The outputs are the estimated cell-type composition matrix 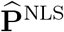, algorithm running time, and list of cell types corresponding to the column name of the resulting matrix.

### Visualization

We provide two functions for the visualization of deconvolution results. The required inputs are ST data and the two outputs of the previous deconvolution step: cell-type composition matrix 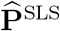 or 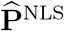 and the list of cell types corresponding to the column name of the resulting matrix. There are optional arguments relating to display, allowing users to adjust their figures flexibly. See details in our document and tutorials.

- *overall plt()* is the command for a spatial scatter pie plot displaying inferred cell-type composition on each location.
- *separate plt()* is the command for spatial proportion plots of given cell types. In Figure 1, we show example visualization results of SLS on the seqFISH+(10000) dataset.

**Figure 1:**
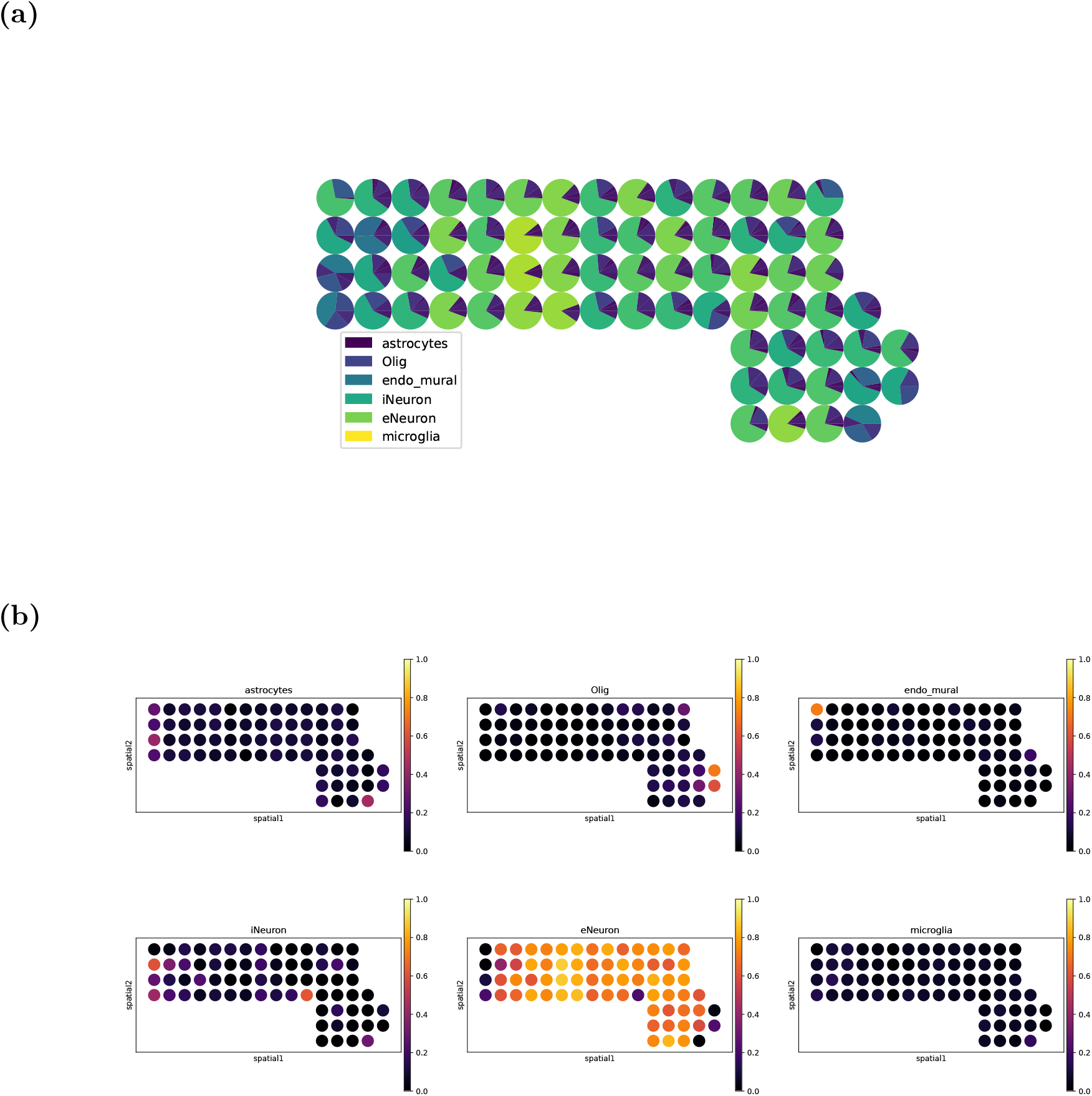
Example visualization results of SLS on the seqFISH+ (10000) dataset. (a) A spatial scatter pie plot displays inferred cell-type composition on spatial location (produced by the command *overall plt()*). (b) The proportion of each cell type is displayed on spatial location (produced by the command *separate plt()*).

## 4 Results

We assess the performance of proposed methods by comparing them with 18 existing methods on diverse benchmarking datasets reported in the recent review paper (Li et al. (2023)). The comprehensive datasets represent two image-based platforms, seqFISH+, MERFISH, and four sequencing-based platforms: Spatial Transcriptomics (ST), 10X Visium (Visium), Slide-seqV2, and stereo-seq. For seqFISH+ and MERFISH datasets, as the gene expression profile is at the single-cell level and cell type labels are known, we follow the exact approach by Li et al. (2023) to bin the neighboring cells into spots with different resolutions and use root-mean-square error (RMSE), Jensen-Shannon Divergence (JSD) to gauge the accuracy of deconvoluted cell type proportion. For the sequencing-based data sets where the true label is unknown, we also follow Li et al. (2023) to use Pearson Correlation Coefficient (PCC) between the deconvoluted cell type proportions within each spot and the expression of the corresponding marker genes of each cell type as the performance metric. Performance of other methods are ported from the Source Data file of Li et al. (2023).

For SLS, it is tuning-free. For NLS, we set *λ* = 0.1 when *n*_*s*_ *> n*_*c*_, and *λ* = 0 when *n*_*s*_ ≤ *n*_*c*_. We set the learning rate to be 0.01, the number of epochs to be 10^3^, and use a warm start. We apply the codes provided by Li et al. (2023) for performance measures to avoid coding bias. Table 1 and 2 show the performance of our methods in comparison to other models on the image-based datasets with true labels. In general, our NLS has the lowest RMSE and JSD, followed by our SLS method, which can be viewed as a fast approximation version of NLS. For the sequencing-based datasets without true labels, to our surprise, the PCC measure suggests that SLS performs even better than NLS, ranking first overall out of all methods (Table 3).

**Table 1:** RMSE of SLS and NLS methods compared with 18 existing methods on image-based datasets seqFISH+ and MERFISH. The numbers in the parenthesis of seqFISH+ indicate the numbers of genes that were randomly chosen in the dataset (We used the fixed random datasets given in Li et al. (2023)); those of MERFISH indicate the binning sizes (in *µ*m) for generating true labels. The last column is the mean of RMSE on all datasets. The top two methods are labeled in bold font. * means equal RMSE when rounding to three decimal places.

|  | seqFISH+ |  |  | MERFISH |  |  | mean |
| --- | --- | --- | --- | --- | --- | --- | --- |
|  | (10000) | (6000) | (3000) | (100) | (50) | (20) |  |
| SLS | **0.109** | 0.113 | **0.126\*** | 0.133 | 0.181 | 0.250 | **0.152** |
| NLS | **0.100** | **0.110** | **0.126\*** | 0.109 | 0.165 | 0.243 | **0.142** |
| Berglund | 0.272 | 0.267 | 0.280 | 0.177 | 0.177 | 0.286 | 0.243 |
| CARD | 0.126 | 0.129 | 0.141 | 0.188 | 0.188 | 0.264 | 0.172 |
| Cell2location | 0.242 | 0.243 | 0.240 | 0.109 | **0.109** | 0.247 | 0.198 |
| DestVI | 0.272 | 0.282 | 0.300 | 0.117 | 0.117 | 0.219 | 0.218 |
| DSTG | 0.227 | 0.260 | 0.292 | 0.231 | 0.231 | 0.311 | 0.259 |
| NMFreg | 0.296 | 0.291 | 0.291 | 0.180 | 0.180 | 0.310 | 0.258 |
| RCTD | 0.125 | 0.130 | 0.131 | 0.184 | 0.184 | 0.278 | 0.172 |
| SD2 | 0.179 | 0.206 | 0.209 | 0.210 | 0.210 | 0.345 | 0.227 |
| SpatialDecon | 0.202 | 0.204 | 0.201 | 0.217 | 0.217 | 0.306 | 0.225 |
| SpatialDWLS | 0.112 | **0.109** | **0.121** | 0.213 | 0.213 | 0.525 | 0.215 |
| SpiceMix | 0.189 | 0.249 | 0.457 | 0.390 | 0.390 | 0.391 | 0.345 |
| SPOTlight | 0.262 | 0.257 | 0.256 | 0.169 | 0.169 | 0.318 | 0.239 |
| STdeconvolve | 0.343 | 0.325 | 0.326 | 0.220 | 0.220 | 0.247 | 0.280 |
| stereoscope | 0.223 | 0.226 | 0.228 | 0.193 | 0.193 | 0.274 | 0.223 |
| STRIDE | 0.260 | 0.273 | 0.232 | 0.124 | 0.124 | 0.372 | 0.231 |
| Tangram | 0.251 | 0.251 | 0.259 | 0.177 | 0.177 | 0.278 | 0.232 |
| SpaOTsc | 0.226 | 0.223 | 0.226 | **0.104** | **0.115** | **0.151** | 0.174 |
| novoSpaRc | 0.227 | 0.227 | 0.225 | **0.096** | 0.124 | **0.162** | 0.177 |

**Table 2:** JSD of SLS and NLS methods compared with 18 existing methods on image-based datasets seqFISH+ and MERFISH. The numbers in the parenthesis of seqFISH+ indicate the numbers of genes that were randomly chosen in the dataset (We used the fixed random datasets given in Li et al. (2023)); those of MERFISH indicate the binning sizes (in *µ*m) for generating true labels. The last column shows the mean JSD on all datasets. The top two methods are labeled in bold font. * means equal RMSE when rounding to three decimal places.

|  | seqFISH+ |  |  | MERFISH |  |  | mean |
| --- | --- | --- | --- | --- | --- | --- | --- |
|  | (10000) | (6000) | (3000) | (100) | (50) | (20) |  |
| SLS | 0.117 | 0.097 | **0.118\*** | 0.093 | 0.185 | 0.296 | **0.151** |
| NLS | **0.090** | **0.086** | **0.118\*** | **0.078** | 0.172 | 0.270 | **0.136** |
| Berglund | 0.380 | 0.373 | 0.425 | 0.169 | 0.169 | 0.172 | 0.281 |
| CARD | 0.142 | 0.140 | 0.157 | 0.225 | 0.225 | 0.319 | 0.201 |
| Cell2location | 0.324 | 0.331 | 0.319 | **0.082** | **0.082** | 0.300 | 0.240 |
| DestVI | 0.443 | 0.474 | 0.519 | 0.089 | 0.089 | **0.172** | 0.298 |
| DSTG | 0.310 | 0.363 | 0.429 | 0.258 | 0.258 | 0.353 | 0.329 |
| NMFreg | 0.489 | 0.478 | 0.479 | 0.185 | 0.185 | 0.457 | 0.379 |
| RCTD | 0.143 | 0.145 | 0.140 | 0.156 | 0.156 | 0.242 | 0.164 |
| SD2 | 0.216 | 0.261 | 0.277 | 0.221 | 0.221 | 0.528 | 0.288 |
| SpatialDecon | 0.275 | 0.279 | 0.270 | 0.236 | 0.236 | 0.282 | 0.263 |
| SpatialDWLS | **0.064** | **0.073** | **0.096** | 0.215 | 0.215 | 1.000 | 0.277 |
| SpiceMix | 0.241 | 0.352 | 0.827 | 0.687 | 0.687 | 0.473 | 0.545 |
| SPOTlight | 0.387 | 0.394 | 0.386 | 0.177 | 0.177 | 0.498 | 0.336 |
| STdeconvolve | 0.544 | 0.477 | 0.516 | 0.267 | 0.267 | **0.065** | 0.356 |
| stereoscope | 0.270 | 0.260 | 0.263 | 0.189 | 0.189 | 0.213 | 0.231 |
| STRIDE | 0.363 | 0.401 | 0.315 | 0.088 | **0.088** | 0.659 | 0.319 |
| Tangram | 0.364 | 0.356 | 0.372 | 0.141 | 0.141 | 0.319 | 0.282 |
| SpaOTsc | 0.648 | 0.638 | 0.654 | 0.167 | 0.268 | 0.462 | 0.473 |
| novoSpaRc | 0.647 | 0.626 | 0.633 | 0.233 | 0.352 | 0.491 | 0.497 |

**Table 3:** Mean PCC of pairs of deconvoluted proportion of cell types and their corresponding marker genes from SLS and NLS methods in comparison with 18 existing methods on sequencing-based datasets ST, Visium, Slide-seqV2, and stereo-seq. The numbers in the parenthesis of Slide-seqV2 indicate the cell-type numbers of the dataset after the sub-cell types are integrated and the original dataset; the names in the parenthesis of stereo-seq indicate the original dataset and integrated subcellular-resolution dataset. The last column is the overall mean of PCC on all datasets. The top two methods are labeled in bold font.

|  | ST | Visium | Slide-seqV2 |  | stereo-seq |  | mean |
| --- | --- | --- | --- | --- | --- | --- | --- |
|  |  |  | (11) | (17) | (gene) | (spot) |  |
| SLS | **0.241** | 0.732 | **0.345** | **0.333** | 0.392 | 0.303 | **0.391** |
| NLS | 0.144 | 0.499 | 0.226 | 0.190 | 0.522 | 0.463 | 0.341 |
| Berglund | 0.063 | 0.553 | 0.195 | 0.195 | 0.520 | 0.026 | 0.259 |
| CARD | 0.117 | 0.730 | 0.250 | 0.200 | 0.233 | **0.471** | 0.333 |
| Cell2location | 0.209 | **0.793** | 0.305 | 0.325 | 0.593 | 0.018 | 0.374 |
| DestVI | 0.081 | 0.642 | -0.055 | -0.070 | 0.537 | 0.449 | 0.264 |
| DSTG | 0.038 | 0.661 | 0.110 | 0.103 | 0.380 | 0.236 | 0.255 |
| NMFreg | -0.103 | 0.496 | 0.190 | 0.243 | 0.533 | 0.400 | 0.293 |
| novoSpaRc | -0.245 | 0.069 | 0.180 | 0.285 | 0.100 | 0.335 | 0.121 |
| RCTD | 0.160 | 0.011 | 0.305 | 0.320 | **0.613** | 0.462 | 0.312 |
| SD2 | 0.033 | 0.585 | 0.075 | 0.100 | 0.410 | 0.149 | 0.225 |
| SpaOTsc | -0.009 | 0.590 | 0.150 | 0.098 | 0.233 | 0.263 | 0.221 |
| SpatialDecon | **0.295** | 0.018 | **0.320** | 0.325 | 0.450 | **0.506** | 0.319 |
| SpatialDWLS | -0.103 | -0.008 | -0.060 | -0.025 | -0.070 | -0.021 | -0.048 |
| SpiceMix | -0.055 | 0.540 | -0.065 | -0.040 | 0.573 | 0.273 | 0.204 |
| SPOTlight | 0.125 | 0.750 | 0.270 | 0.233 | 0.360 | 0.433 | 0.362 |
| STdeconvolve | -0.030 | **0.760** | 0.300 | 0.213 | **0.603** | 0.133 | 0.330 |
| stereoscope | 0.146 | 0.668 | 0.260 | 0.255 | 0.527 | 0.436 | 0.382 |
| STRIDE | -0.062 | 0.642 | 0.235 | 0.205 | 0.543 | 0.434 | 0.333 |
| Tangram | 0.071 | 0.743 | 0.290 | **0.365** | 0.510 | 0.360 | **0.390** |

One appealing feature of the SLS method is its computing efficiency. For a comparison, we pick three methods that were shown relatively more efficient in Li et al. (2023), including Tangram (Biancalani et al., 2021), RCTD (Cable et al., 2022) and SpatialDecon (Danaher et al., 2022). We consider benchmarking on two example datasets: seqFISH+ (10000), which represents datasets with large gene numbers and small spot numbers, and MERFISH (20), which represents datasets with large spot numbers and small gene numbers. We run SLS and the other three methods on a cluster node with Intel(R) Xeon(R) Gold 6338 CPU @ 2.0GHz and 256GB DDR4 2666MHz Memory with allocation of 1 CPU core and 100GB memory. We run NLS on Google Colab T4 GPU. The computing time is shown in Table 4. We note that Tangram is also computationally efficient. Nevertheless, its performance is less competitive. In conclusion, we recommend SLS for users with limited computation resources and NLS for those with high accuracy standards and a demand for code flexibility.

**Table 4:**
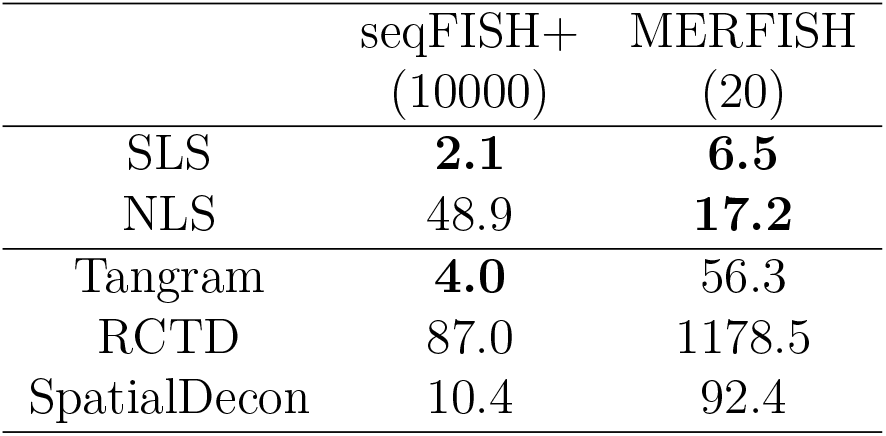
Running time (seconds) on seqFISH+(10000) and MERFISH(20) datasets. We show the top two methods in bold.

## Acknowledgments

Yunlu Chen thanks Hangyu Lin and Chenhao Zhang for helpful discussion on this project.

